# MSC-delivered High-Affinity Variants of soluble PD1 lead to tumour regression

**DOI:** 10.64898/2025.12.04.692338

**Authors:** Serap Gokcen, Tianyuan Chu, Phoebe Blair, Natalia Krajewska, Greg Brooke, Stuart Rushworth, Ralf Michael Zwacka, Andrea Mohr

## Abstract

Immune checkpoint therapies aim to restore anti-tumour immunity by blocking inhibitory signals that suppress T-cell activation. The most effective current strategies use humanised antibodies targeting PD1 or its ligand PDL1. However, due to their large size, antibodies often exhibit limited tissue diffusion, resulting in poor penetration into solid tumours. To address this challenge, we developed a novel checkpoint inhibitor approach that uses mesenchymal stromal cells (MSCs) to deliver a high-affinity, soluble PD1 receptor (sPD1HAC). We found that sPD1HAC produced as an IgG1-Fc fusion protein provided functional expression in MSCs. The sPD1HAC-IgG1-Fc fusion protein showed strong and specific binding to PDL1 and could outcompete recombinant PD1 and anti-PDL1 antibodies, including the clinically approved durvalumab. Although the sPD1HAC variant was designed to block human PD1-PDL1 signalling, it also bound murine PDL1 and blocked the binding of mouse-specific PD1 antibodies as well as recombinant murine PD1 protein. Accordingly, we found significant anti-tumour activity of intravenously administered MSCs.PD1^HAC^ in an aggressive B16-F10 cancer model. This cell-based immune checkpoint approach offers a potential therapeutic option for targeting stroma-rich, treatment-resistant tumours.

## Introduction

Immune checkpoint therapies have become a game-changer in the treatment of several cancer types, including melanoma, non-small cell lung cancer, and renal cell carcinoma (1, 2). Immune checkpoint molecules, such as cytotoxic T-lymphocyte-associated protein 4 (CTLA-4) (3), programmed cell death protein 1 (PD1) (4), and its ligand PDL1 (5, 6), are essential regulators of immune tolerance, preventing autoimmunity by modulating T-cell activity (7, 8). However, tumours can hijack these pathways to evade immune surveillance, promoting immune tolerance and facilitating cancer progression (9). Therefore, these immune checkpoints have become key therapeutic targets and immune checkpoint inhibitors (ICIs) have been successfully developed and used clinically. ICIs are typically monoclonal antibodies designed to block the interaction between the ligand and receptor, thereby restoring effective T-cell-mediated anti-tumour responses (10). Since the approval of ipilimumab, the first anti-CTLA-4 ICI, in 2011, several agents targeting the PD1/PDL1 axis, such as nivolumab and pembrolizumab (anti-PD1), and atezolizumab and durvalumab (anti-PDL1), have been added to the arsenal, and all have shown remarkable clinical efficacy (11, 12).

The PD1/PDL1 pathway plays a particularly central role in immune checkpoint regulation. PD1, encoded by the *PDCD1* gene, is a type I transmembrane protein expressed predominantly on activated T cells, B cells, and myeloid cells. Structurally, PD1 contains an immunoglobulin V-like extracellular domain, a transmembrane region, and a cytoplasmic tail with two key signalling motifs: an immunoreceptor tyrosine-based inhibitory motif (ITIM) and an immunoreceptor tyrosine-based switch motif (ITSM). Upon engagement with its ligands, PDL1 (B7-H1) and PDL2 (B7-DC), PD1 undergoes phosphorylation at these motifs, initiating downstream inhibitory signalling cascades. These phosphorylation events lead to the recruitment of the protein tyrosine phosphatase SHP-2 (13), impairing the activation of downstream pathways critical for T-cell activation and proliferation (14, 15). Along with these effects, PD1 signalling inhibits the activation of transcription factors, including AP-1 and NFAT, which are crucial regulators of cytokine production and T-cell differentiation. The impact on NF-κB is more nuanced, suggesting a complex interplay rather than simple inhibition (16, 17). Together, this promotes T-cell exhaustion, a hyporesponsive state characterised by reduced effector function and sustained expression of inhibitory receptors (18).

Of the two ligands, PDL1 and PDL2, PDL1 plays the more dominant role in tumour immune evasion (19). It is prominently found on dendritic cells, macrophages, monocytes, B cells, and activated T-cells. Its expression on dendritic cells and macrophages is vital for modulating T-cell activation and maintaining peripheral tolerance. Myeloid-derived suppressor cells (MDSCs) also exhibit high levels of PDL1, which contributes to their immunosuppressive function in pathological contexts like cancer (20). In the non-immune compartment, PDL1 is expressed on epithelial and endothelial cells, particularly under inflammatory conditions mediated by cytokines such as interferon-γ (IFN-γ), tumour necrosis factor-α, and interleukin-6 (21–24). Cancer-associated fibroblasts (CAFs) within the tumour microenvironment (TME) can also express PDL1, further reinforcing local immune suppression (25). Most notably, PDL1 is frequently upregulated on tumour cells across a broad spectrum of malignancies, enabling evasion of cytotoxic T lymphocyte-mediated killing (26).

Blocking the PD1/PDL1 interaction with ICIs such as pembrolizumab or durvalumab can reactivate T cells and restore anti-tumour immunity. However, treatments with blocking antibodies are associated with several issues, so-called immune-related adverse events (irAEs), where the immune system attacks healthy tissues (27, 28). The two most frequent effects are dermatologic toxicities, such as rash, itchy skin, and vitiligo, and gastrointestinal toxicities, including diarrhoea and colitis (29, 30), which could potentially be overcome by cell-therapeutic delivery of ICIs. In this context, mesenchymal stem/stromal cells (MSCs) offer a variety of advantages. MSCs can be engineered to act as “Trojan horses,” delivering therapeutic proteins directly to tumour sites, owing to their inherent ability to migrate to tumour tissues, thereby improving targeting and potentially reducing systemic toxicity (31, 32). The natural ability of MSCs to migrate toward tumour tissues enhances their utility in overcoming the poor tissue penetration that often limits the effectiveness of monoclonal antibodies in solid tumours. Unlike passive antibody therapies with limited half-lives, MSCs can persist, expand, and adapt within the TME, enabling long-term and continuous therapeutic action (33–35).

We generated MSCs that express and secrete a soluble PD1 (sPD1) ectodomain, which acts as a competitive inhibitor of PDL1-PD1 binding. To produce a high-affinity PD1 variant, ten specific point mutations were introduced into the sPD1 sequence. This so-called HAC variant was first described by Maute et al. (2016) and has shown a significantly increased binding affinity to PD-L1 (36). We analysed this variant and wild-type sPD1 (sPD1^WT^) in several *in vitro* and *in vivo* systems, including MSC-delivery in a melanoma model.

## Materials and Methods

### Cell Culture

The following human cancer cell lines were obtained from the American Type Culture Collection (ATCC, Manassas, VA, USA): BxPC3, Colo-357, HCT116, LoVo, RKO, MDA-MB-231, T-47D, MCF-7, PC-3, Du145, LNCaP, HeLa, A549, A2780, CHO, HEK293T, LL/2-Luc, MC38, and B16-F10. PancTu1 cells were kindly provided by the Kalthoff laboratory. MM.1R, MM.1S, U266, and bone marrow-derived MSCs (BM-MSCs) were purchased from Cytion Biosciences (Eppelheim, Germany), while umbilical cord-derived MSCs (UC-MSCs) were obtained from PromoCell (Heidelberg, Germany). 5TGM1 cells were generously provided by Dr Stuart Rushworth’s laboratory, and adipose tissue-derived MSCs (AT-MSCs) were purchased from AMSBio (Abingdon, United Kingdom). CHO, HEK293T, MCF-7, MDA-MB-231, LoVo, Du145, HeLa, A549, LL/2-Luc, MC38, and B16-F10 cells were cultured in Dulbecco’s Modified Eagle Medium (DMEM; Corning Inc., Corning, NY, USA). PancTu1, BxPC3, Colo-357, LNCaP, PC-3, MM.1R, MM.1S, U266, T-47D, A2780, and 5TGM1 cells were maintained in Roswell Park Memorial Institute 1640 medium (RPMI-1640; Corning Inc.). RKO and HCT116 cells were grown in McCoy’s 5A Modified Medium (Corning Inc.), and all MSCs were cultured in MesenPRO RS medium (Thermo Fisher Scientific, Waltham, MA, USA). All media were supplemented with 10% fetal bovine serum (FBS; Thermo Fisher Scientific), 100 IU/mL penicillin, and 100 μg/mL streptomycin (both from Thermo Fisher Scientific). LL/2-Luc cells were additionally supplemented with 2 μg/mL blasticidin (Thermo Fisher Scientific). Cells were maintained at 37 °C in a humidified incubator with 5% CO₂ and passaged using trypsin-EDTA (Lonza, Basel, Switzerland).

### Reagents including antibodies

PE-conjugated anti-human PDL1 and PD1 antibodies (referred to as αhPDL1 and αhPD1, respectively; BioLegend, San Diego, CA, USA; Cat. Nos. 329705 and 329905) were used for PDL1 and PD1 detection on human cells, as well as in antibody binding competition assays. For murine samples, PE-conjugated anti-mouse PDL1 antibodies, MIH7, MIH6, and 10F.9G2 (BioLegend; Cat. Nos. 155403, 153611, and 124301, respectively) were employed for PDL1 detection and competition assays. These antibodies are referred to by their clone names or collectively as αmPDL1. Additionally, a PE-conjugated durvalumab biosimilar monoclonal antibody (Cambridge Bioscience, United Kingdom; Cat. No. BME100153P), referred to as DUR (PE), was used as a probe in antibody binding competition assays. Isotype controls included PE-conjugated mouse IgG2bκ (BioLegend; Cat. No. 400311) for αhPDL1/αhPD1, PE - conjugated rat IgG2bκ (BioLegend; Cat. No. 400607) for 10F.9G2, PE-conjugated rat IgG2aκ (BioLegend; Cat. No. 400508) for MIH6 and MIH7, and PE-conjugated human IgG1κ (BD Biolegend; Cat. No. 403503) for DUR (PE). In both solid-phase and cell-based competition assays, biotin-tagged recombinant human PD1 Fc-fusion protein (rhPD1; BioLegend; Cat. No. 799506) was used as probe. Unconjugated durvalumab (Selleck Chemicals; Houston, TX, USA Cat. No. A2013) was included as a competitor alongside sPD1 variants to evaluate blocking efficiency against rhPD1. For solid-phase competition assays, recombinant human PDL1 Fc chimaera protein (rhPDL1; Biotechne (Minneapolis, MN, USA); Cat. No. 156-B7-100) was used as the coating protein. Gemcitabine (Cat. No. S1149) was purchased from Selleck Chemicals and interferon-γ from Biotechne (Cat. No. 285-IF-100/CF).

### Generation of sPD1 and PDL1 constructs

sPD1**^WT^** corresponds to the extracellular domain of human PD1 (amino acids 25–167), while sPD1**^HAC^** represented a high-affinity consensus variant containing 10 targeted point mutations, as previously described (36). Both constructs were cloned into the pFUSE-hIgG1-Fc vector (InvivoGen, San Diego, CA, United States), downstream of a human fibrillin-1-derived signal peptide and a furin cleavage site to promote PD1 secretion (31, 37, 38). To assess secretion efficiency and protein stability, constructs were designed either with (stWT, stHAC) or without (WT, HAC) a stop codon between the PD1 fragment and the hIgG1-Fc domain. The murine PDL1 construct (mPDL1) was purchased from Genescript (clone ID OMu23190; Piscataway, NJ, USA). Adenoviral vectors encoding sPD1**^HAC^** (Ad.sPD1**^HAC^**) and a luciferase control (Ad.LUC) were obtained from Vector Biolabs (Philadelphia, PA, USA).

### Cell transfections and transductions

CHO and HEK293T cells (1 × 10⁶ cells per well) were seeded in 6-well plates and transfected using TurboFect transfection reagent (Thermo Fisher Scientific) following the manufacturer’s protocol, with a reagent-to-DNA ratio of 3:1. For sPD1 expression, cell culture supernatants were collected 48 h post-transfection. For PDL1 expression analysis, HEK293T cells were harvested 24 h post-transfection and assessed for surface ligand expression by flow cytometry. MSCs were transduced with E1/E3-deleted adenoviral vectors expressing either sPD1^HAC^ (Ad.CMV.sPD1^HAC^) or luciferase (Ad.CMV.LUC, control) at varying multiplicities of infection (MOIs). MSCs were transduced as previously described (39).

### Detection of sPD1 Secretion by Enzyme-Linked Immunosorbent Assay (ELISA)

sPD1 protein secretion in the culture supernatants of transfected CHO and HEK293T cells, as well as transduced MSCs, was quantified using the human PD1 DuoSet ELISA kit (Biotechne; Cat. No. DY1086), in accordance with the manufacturer’s instructions. Briefly, cell supernatants were collected 48 h post-transfection or post-transduction, cleared by centrifugation to remove cellular debris, and appropriately diluted prior to PD1 quantification by ELISA.

### Cell Surface Staining and FACS Analysis

Surface expression was assessed by incubating cells with directly conjugated antibodies, using a staining protocol previously described (38). Matched isotype control antibodies were included to evaluate non-specific binding. Following staining, cells were washed with phosphate-buffered saline (PBS), fixed in 4% paraformaldehyde (PFA), and analysed by flow cytometry.

### Solid-phase binding competition assay

96-well plates were coated overnight at 4 °C with recombinant human PDL1 (rhPDL1; 50 ng/well), followed by three washes with PBS. Wells were then blocked with dilution reagent for 1 h at room temperature. Subsequently, plates were incubated for 2 h with a mixture of biotinylated rhPD1 (50 ng/well) and serial dilutions of sPD1 variant-containing supernatants (ranging from 40 to 0.01 ng/well) or molar-equivalent concentrations of durvalumab in PBS. Control wells contained biotinylated rhPD1 and cell supernatant from untransfected cells. After incubation, wells were washed and re-blocked for 10 min. Streptavidin–horseradish peroxidase (HRP; 1:40 dilution; Bio-Techne) was added for 20 min, followed by colourimetric detection using Tetramethylbenzidine (TMB) substrate. Optical density (OD) was measured at 450 nm, and rhPD1 binding (%) was calculated using the formula: (OD sample / OD control) × 100.

### Cell-based binding competition assay

RKO cells (4 × 10⁵) were incubated on ice for 25 min with a mixture containing biotinylated rhPD1 (used as the primary binding probe) and a competitor. Competitors included either sPD1 variants (sPD1^WT^) secreted by HEK293T, CHO, or MSCs or a molar-equivalent concentration of durvalumab. Following incubation, cells were washed and stained with Streptavidin-PE (1:25 dilution in PBS) for 20 min at 4 °C, fixed in 4% PFA, and analysed by flow cytometry.

To normalise the data, background fluorescence, measured from cells treated with PBS and Streptavidin-PE only, was subtracted from all samples. Corrected rhPD1 binding in the absence of competitor was defined as 100%. The percentage of rhPD1 binding in the presence of each competitor was calculated using the formula:

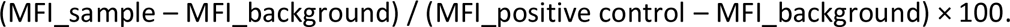

### Antibody binding competition assay

Cells (4 × 10⁵) were incubated for 25 min on ice with a mixture of PE-conjugated αPDL1 antibody (including a DUR (PE), used as binding probe, and either sPD1**^HAC^**, sPD1**^WT^**or MSC.LUC supernatants as competitors. The sPD1 variants were applied at 0.8 μg per sample. Controls consisted of cells incubated with αPDL1 antibody alone (either mixed with PBS or MSC.LUC supernatant), while negative controls were incubated with the corresponding isotype control. Following incubation, cells were washed with PBS, fixed in 4% PFA, and analysed by flow cytometry. MFI values were normalised and calculated as described for the cell-based binding competition assays.

### Tumour model

C57BL/6 mice were housed under specific pathogen-free conditions in a Containment Level 3 disease modelling unit. All procedures were conducted in accordance with the UK Animals (Scientific Procedures) Act 1986 (revised 2013) and approved by the Home Office. On day 0, pulmonary metastases were established by tail vein injection of B16.F10 melanoma cells (2 × 10⁵ cells in 100 µL PBS). On day 7, mice received a single intravenous dose of either MSC.sPD1^HAC^ (4 × 10⁵ cells in 100 µL PBS) or MSC.LUC as a control. Animals were euthanised 7 days post-treatment for endpoint analyses. For histopathological analysis, the lungs were fixed in 10% neutral buffered formalin, embedded in paraffin, sectioned at a thickness of 4 µm, and stained with Haematoxylin and Eosin (H&E). Sections were examined under a light microscope to assess the presence and extent of metastatic lesions.

### DNA-hypodiploidy assays

Apoptosis was measured according to Nicoletti et al. (32). Briefly, 24 h post-transfection/transduction, cells were harvested. Pelleted cells were resuspended in hypotonic fluorochrome solution containing 20 μg/ml propidium iodide, 0.1% (w/v) sodium citrate, and 0.1% (v/v) Triton-X100. After incubation at 4 °C for 20 min, cells were analysed by flow cytometry. Cell death was quantified as the proportion of cells in the sub-G1 phase, indicative of DNA fragmentation. Viable cell percentage was calculated by subtracting the proportion of dead cells from the total population (100%).

### Cytokine array

Cellular supernatants were harvested from MSCs or MSC-LUC 48 h after transduction. The supernatants were centrifuged at 13,300 rpm for 15 min to remove cellular debris. After a five-fold dilution, the supernatants were applied to Human Cytokine Antibody Array C3 membranes (RayBiotech), according to the manufacturer’s instructions. The final images from the CCD camera were analysed by subtracting the background signal and normalising each cytokine signal to the internal positive control.

### Statistical Analysis

Experimental values are expressed as mean value±standard error of the mean (SE). For significance analyses, analysis of variance (ANOVA) was used, and p ≤ 0.05 was considered significant. Experiments were performed three times unless otherwise stated.

## Results

### Fusion of sPD1 to an IgG1-Fc Domain Enhances Protein Stability and Secretion

The PD1 receptor is a type I membrane protein of 288 amino acids (Figure 1A). We used the ectodomain of PD1 to generate sPD1-WT and the sPD1-HAC variant (Figure 1A) that were expressed as either IgG1-Fc fusion or non-fusion proteins (Figure 1B). The successful secretion of these proteins from producer cells, such as HEK293T or CHO cells, was mediated by the signal peptide from the human Fibrillin protein (aa 1-aa 27), and a Furin cleavage site (Figure 1A-B).

**Figure 1.**
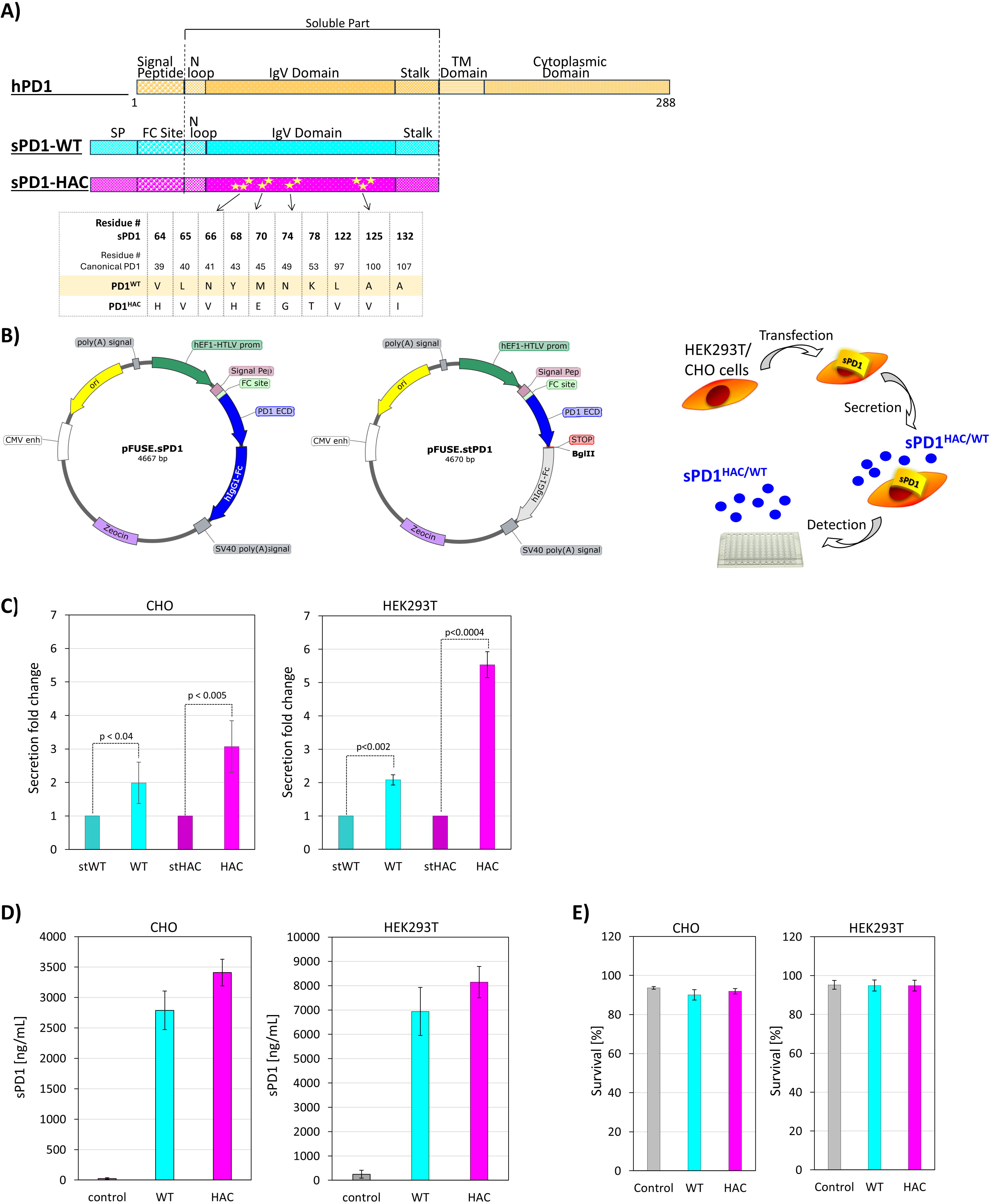
Generation, secretion, and comparative analysis of sPD1-WT and sPD1-HAC fusion and non-fusion proteins. **A** Overview of the sPD1-WT (turquoise) and sPD1-HAC (pink) constructs, which consist of the ectodomain of the membrane-bound full-length PD1 (orange). The 10-point mutations introduced to generate the high-affinity HAC variant are shown in the sPD1-HAC construct. **B** HEK293T and CHO cells were transfected with the depicted constructs to produce and secrete IgG1-Fc-sPD1-WT and IgG1-Fc-sPD1-HAC fusion proteins, as well as sPD1-WT and sPD1-HAC proteins. For these, a stop codon was inserted at the 3’-end of the sPD1-WT and sPD1-HAC sequences within the respective IgG1-Fc constructs, based on the pFUSE-hIgG1-Fc1 plasmid. The addition of the stop codon allowed for direct comparison of the fusion and non-fusion proteins in terms of yield and function. **C** The concentrations of sPD1-WT (stWT), IgG1-Fc-sPD1-WT (WT), sPD1-HAC (stHAC), and IgG1-Fc-sPD1-HAC (HAC) proteins in supernatants of transfected CHO and HEK293T cells were analysed by PD1 ELISA. The expression of sPD1-WT (stWT) and sPD1-HAC (stHAC) was set to 1. Changes in expression relative to stWT and stHAC are presented as fold changes. Error bars represent mean values ± SE. **D** A comparison of protein expression levels of IgG1-Fc-sPD1-WT (WT) and IgG1-Fc-sPD1-HAC (HAC) in the supernatants of transfected CHO and HEK293T cells. Protein concentrations, expressed in ng/ml, were measured by ELISA. Supernatants from mock-transfected cells (control) served as control. **E** The viability of IgG1-Fc-sPD1-WT (WT) or IgG1-Fc-sPD1-HAC (HAC) expressing CHO and HEK293T cells was assessed using a DNA hypodiploidy assay and expressed as cell survival (%) values. All results are expressed as mean values ± SE. The constructs and proteins designated IgG1-Fc-sPD1-WT and IgG1-Fc-sPD1-HAC will be referred to as sPD1^WT^ and sPD1^HAC^, respectively, in all subsequent experiments and figures.

When we compared the expression and secretion of the different sPD1 variants, we found that IgG1-Fc fusion proteins yielded significantly higher levels in the supernatant of the two producer cell lines than their non-fusion counterparts (Figure 1C). In particular, the IgG1-Fc-sPD1-HAC fusion protein showed high levels of expression and stability The concentration of IgG1-Fc-sPD1-WT was measured at 2800 ng/mL in CHO cells, whereas the IgG1-Fc-sPD1-HAC construct resulted in 3400 ng/mL in the same cells. In HEK293T cells, the average end concentrations 48 h after transfection were 7000 ng/mL and 8000 ng/mL, respectively (Figure 1D). The expression of these sPD1 proteins did not affect the survival of the producing CHO and HEK293T cells (Figure 1E), demonstrating that cellular delivery of sPD1 variants as ICIs is feasible. As for the higher expression levels of the IgG1-Fc fusion proteins, we used them in all subsequent experiments, referring to them as sPD1^WT^ and sPD1^HAC^ from here on.

### The sPD1^HAC^ outcompetes rhPD1 in ligand binding

In a solid-phase, cell-free competition assay, we examined whether sPD1^WT^ and sPD1^HAC^, produced in CHO and HEK293T cells, could compete with rhPD1 for binding to immobilised rhPDL1. In the assay, the rhPD1 was biotin-tagged and binding to rhPDL1 quantified via Streptavidin-HRP and TMB staining (Figure 2A). The rhPD1 (50 ng) was incubated together with different amounts (40 ng – 0.1 ng) of sPD1^WT^ or sPD1^HAC^ (Figure 2B). The results reveal that only sPD1^HAC^ but not sPD1^WT^ could successfully compete with rhPD1 (Figure 2B). The results demonstrated that even at a 100-fold excess, sPD1HAC could block almost all rhPD1 binding to PDL1. When rhPDL2 was used as a coating agent sPD1^HAC^ lost its inhibitory activity showing that its binding is highly specific for PDL1 and will not block the PD1-PDL2 binding. (Figure 2C).

**Figure 2.**
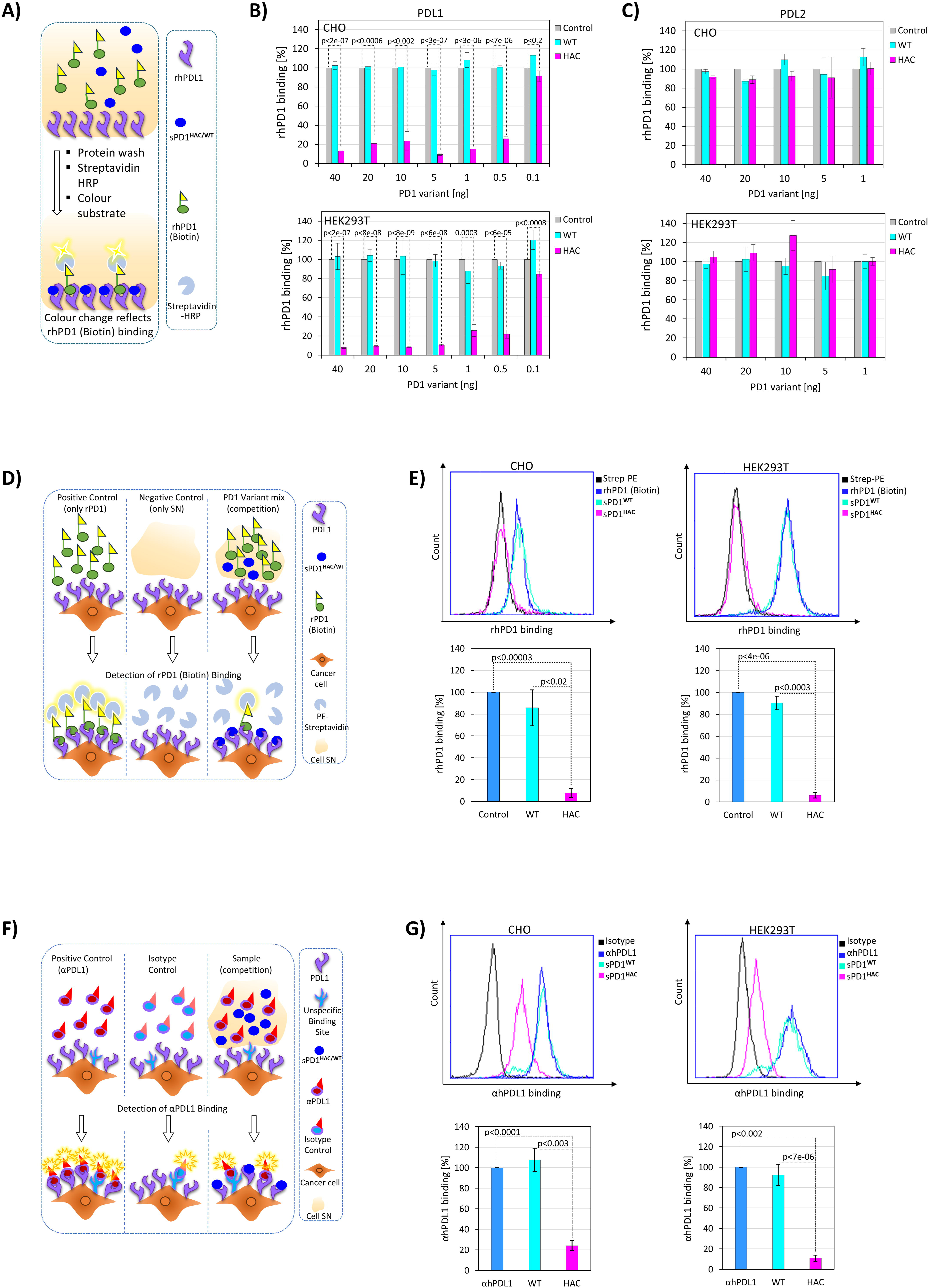
sPD1HAC specifically binds to PDL1 and outcompetes rhPD1 and anti-PDL1 antibodies for binding. **A** Schematic depiction of the experimental setup where wells of a 96-well plate were coated with rhPDL1. The wells were then incubated with rhPD1-Biotin in the presence or absence of sPD1-^HAC/WT^, followed by Streptavidin-HRP and an HRP substrate. A decrease in signal indicates that sPD1^HAC/WT^ outcompeted the biotinylated rhPD1. **B** To assess the ability of sPD1^WT^ (turquoise) and sPD1^HAC^ (pink), produced in CHO (upper panel) or HEK293T (lower panel) cells, to compete for binding to rhPDL1, various amounts (0.1 to 40 ng) of sPD1^WT^ and sPD1^HAC^ were added to the mixture. As controls, equivalent volumes of supernatants from mock-transfected cells were used (grey). **C** The wells of a 96-well plate were coated with rhPDL2. Experimental conditions and setup were identical to panel B. **D** A schematic depiction of the experimental setup where RKO cells were incubated with rhPD1-Biotin, in the presence or absence of sPD1-^HAC/WT^, followed by Streptavidin-PE. A decrease in the PE-signal indicates that sPD1^HAC/WT^ outcompeted the biotinylated rhPD1. **E** Flowcytometric analyses (left) of RKO cells stained only with Streptavidin-PE (negative, unstained control; black), and cells stained with rhPD1-Biotin and Streptavidin-PE plus either CHO (left) or HEK293T (right) cell-produced sPD1^HAC^ (pink), sPD1^WT^ (turquoise) or an equivalent volume from the respective mock-transfected producer cells (blue). Representative FACS diagrams are shown on the top, and quantitative analyses are displayed at the bottom. The results are the mean ± SE. **F** A schematic depiction of the experimental setup where RKO cells were incubated with a PE-conjugated anti-PDL1 antibody, in the presence or absence of sPD1-^HAC/WT^, followed by Streptavidin-PE. A decrease in the PE-signal indicates that sPD1^HAC/WT^ outcompeted the PDL1-specific antibody. **G** Flowcytometric analyses (left) of RKO cells stained only with a PE-conjugated isotype control (black), and cells stained with anti-PDL1-PE plus either CHO (left) or HEK293T (right) cell-produced sPD1^HAC^ (pink), sPD1^WT^ (turquoise) or an equivalent volume from the respective mock-transfected producer cells (blue). Representative FACS diagrams are shown on the top, and quantitative analyses are displayed at the bottom. The results are the mean ± SE.

To ensure sPD1^HAC^ exhibits the same binding affinity for endogenous membrane-bound PDL1 as for recombinant plate-bound PDL1, binding and competition were tested in cellular assays. For this, we tested an array of different cancer cell lines from various cancer types for their PDL1 (and PD1) surface expression. We found that RKO colorectal cancer cells and MDA-MB-231 breast cancer cells exhibited robust PDL1 expression, whereas many others showed little or no endogenous PDL1 expression (Supplemental Figure S1). Only when treated with IFN-γ the majority of cells tested, started to produce PDL1 (Supplemental Figure S1). Based on these results, we decided to use unstimulated RKO cells as a cellular model in the subsequent experiments. When we labelled RKO cells with rhPD1-Biotin and visualised binding with Streptavidin-PE and flow cytometry (Figure 2D), we could readily detect PDL1 expression on the cells (Figure 2E). While the addition of sPD1^WT^ expressed by HEK293T or CHO cells could not interfere with this binding, sPD1^HAC^ reduced the fluorescence signal to near control levels (Figure 2E). Similarly, when we used an anti-PDL1 antibody (Figure 2F), sPD1^HAC^ could significantly block antibody binding, whereas sPD1^WT^ could not (Figure 2G). Taken together, we could show that sPD1^HAC^ could outcompete rhPD1 as well as a specific anti-PDL1 antibody, demonstrating that it is a highly potent, cell-expressed inhibitor of PD1-PDL1 binding.

### MSCs afford robust production and secretion of sPD1^HAC^

To identify the best type of MSCs for our cell therapeutic approach, we tested several MSC types for PDL1 (and PD1) expression and found that adipose tissue-derived MSCs (AT-MSCs) do not express PDL1 (Figure 3A). This is in contrast to bone-marrow-derived MSCs (BM-MSCs) and umbilical cord-derived MSCs (UC-MSCs) that display substantial levels of PDL1. This makes AT-MSCs the better cellular delivery vehicle in this context, as they will not bind and scavenge sPD1^HAC^ themselves, thereby avoiding dilution of the therapeutic effect. In order to engineer MSCs that produce and secrete sPD1^HAC^ (MSC.sPD1^HAC^), the underlying expression cassette was cloned into an adenoviral vector termed Ad.CMV.sPD1^HAC^ (Figure 3B). Subsequently, we used Ad.CMV.sPD1^HAC^ to transduce AT-MSCs (from here on referred to as MSCs) at different MOIs (Figure 3C). After 48 h, the supernatants were harvested and analysed by ELISA for sPD1^HAC^ expression. The results revealed that sPD1^HAC^ was successfully produced and secreted by MSCs in an MOI-dependent manner, with the maximum concentration reaching levels of 800 ng/mL of sPD1^HAC^ (Figure 3C). The transduction with Ad.CMV.sPD1^HAC^ did not affect cell viability compared to non-transduced control MSCs (Figure 3D). In addition, production of sPD1^HAC^ did not alter the cytokine secretion of MSCs, indicating that they will maintain their normal biological properties, including their tissue/tumour tropism (Figure 3E). Next, we tested the ability of MSC-produced sPD1^HAC^ to block binding of rhPD1 to rhPDL1 in a cell-free assay and found that as little as 0.6 ng sPD1^HAC^ had a significant inhibitory effect (Figure 3F). On RKO cells, MSC-produced sPD1^HAC^ could fully compete with the binding of rhPD1 (Figure 3G) as well as a specific anti-PDL1 antibody (Figure 3H) to endogenous PDL1 on the surface of the cells. These results demonstrate that MSCs can be successfully engineered to express and secrete functional sPD1^HAC^ protein.

**Figure 3.**
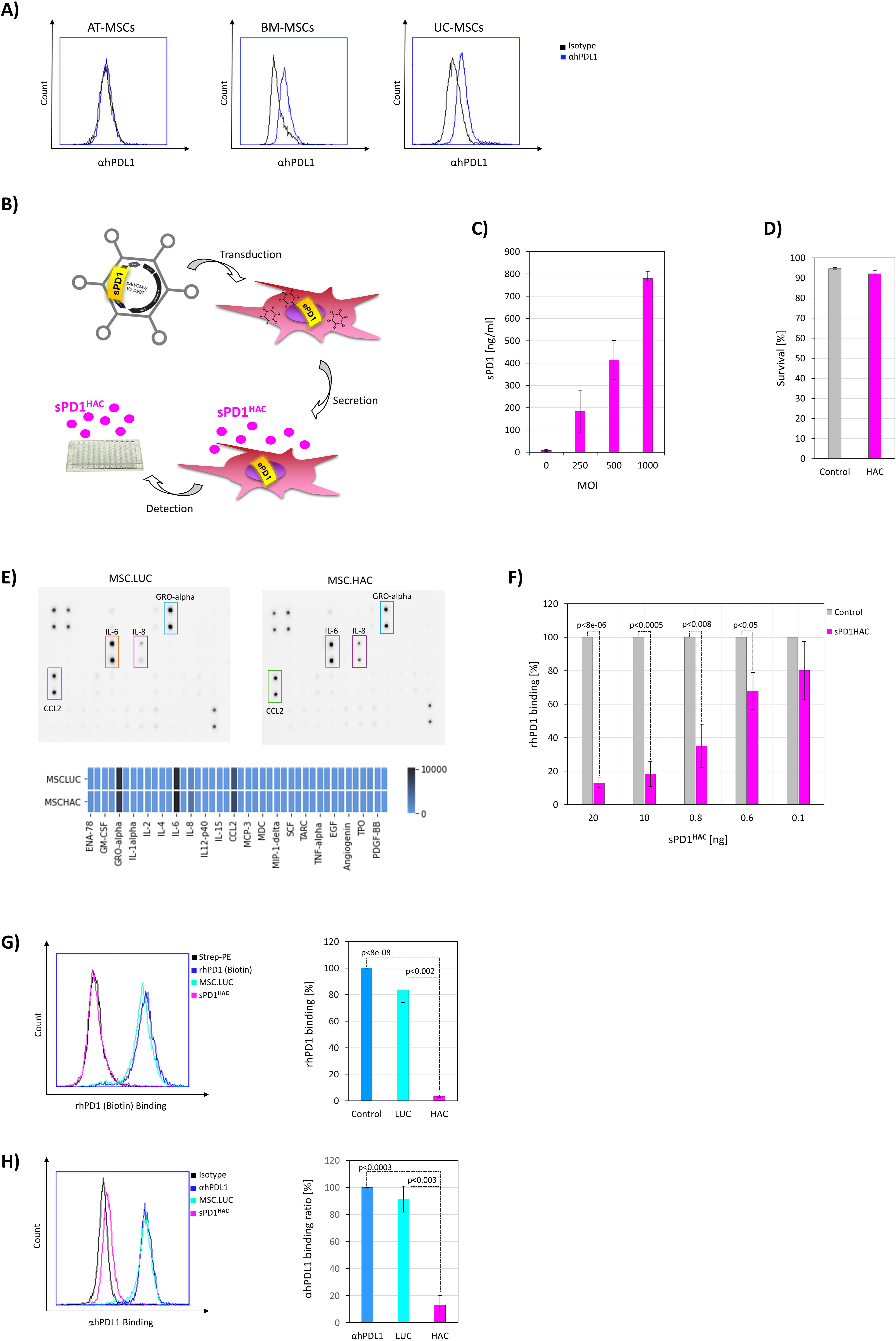
MSC-derived sPD1HAC outcompetes rhPD1 and antibody binding to PDL1 on cancer cells. **A** Three different types of MSCs (AT-MSCs, BM-MSCs and UC-MSCs) were tested for their PDL1 (blue) expression levels by flow cytometry. Isotype controls are shown in black. **B** Schematic representation of the engineering of MSCs to produce and secrete sPD1^HAC^ (MSC.sPD1HAC). The sPD1HAC expression cassette was cloned into an adenoviral vector (Ad.CMV.sPD1HAC), which was subsequently used to transduce MSCs for the secretion of sPD1HAC. **C** PD1 ELISA of MSCs transduced with increasing MOIs (0–1000) of Ad.CMV.sPD1HAC. **D** The impact of sPD1HAC expression on MSCs was assessed using a DNA hypodiploidy assay. Cell survival was calculated by subtracting the percentage of cells showing DNA fragmentation from the total population (100%). The survival of MSCs PD1HAC cells transduced at an MOI of 1000 is shown in pink, with non-transduced MSCs (grey) serving as controls. **E** Cytokine/growth factor array analysis and heatmap visualisation of supernatants from MSCs transduced with a control vector (Ad.LUC), resulting in MSC.LUC, and supernatants from MSC.sPD1^HAC^ (MSC.HAC). Medium supernatants were collected 48 h post-transduction. Factors that are highly expressed in MSCs are highlighted; however, no differences between MSC.LUC and MSC.HAC were observed. **F** MSC-produced sPD1^HAC^ (0.1–20 ng, pink) was added to wells coated with rhPDL1, followed by incubation with rhPD1 Biotin, Streptavidin-HRP, and visualisation by HRP substrate. Signal levels obtained with equal volumes of supernatants from non-transduced control MSCs (grey) were set to 100. G Flow cytometric analysis of RKO cells stained with rhPD1Biotin and Streptavidin-PE (blue), Streptavidin-PE-alone as a negative control (black), or rhPD1Biotin and Streptavidin-PE in the presence of either sPD1^HAC^ (pink) or an equivalent volume of MSC.LUC supernatant (turquoise). Representative FACS plots are shown on the left, with quantitative analyses on the right. Data represent-the mean ± SE of three independent experiments. **H** Flow cytometric analysis of RKO cells stained with antihPDL1-PE antibody (blue), PE-conjugated isotype control antibody (black), or antihPDL1-PE antibody in the presence of either sPD1^HAC^ (pink) or an equivalent volume of MSC.LUC supernatant (turquoise). Representative FACS plots are shown on the left, with quantitative analyses on the right. Data represent mean ± SE.

### The sPD1^HAC^ variant demonstrates enhanced efficacy over durvalumab

To further evaluate sPD1^HAC^, we tested its biological activity against the established ICI, durvalumab (Imfinzi). durvalumab is a human immunoglobulin G1 kappa monoclonal antibody that recognises PDL1 and blocks its interaction with PD1. It is used in the treatment of various cancers, including non-small cell lung cancer (NSCLC), small cell lung cancer (SCLC), bladder cancer, biliary tract cancer, and endometrial cancer, as well as urothelial and hepatocellular carcinoma (40–46). When we compared the activity of durvalumab and sPD1^HAC^ in competing with rhPD1 for binding to immobilised rhPDL1, we observed that sPD1^HAC^ performed equally well as durvalumab and even better at lower concentrations (Figure 4A). Furthermore, when we tested durvalumab and sPD1^HAC^ at equimolar amounts for their potential to compete with rhPD1 on RKO cells, we found that sPD1^HAC^ had higher inhibitory activity (Figure 4B). We also analysed the ability of HEK293T- and MSC-produced sPD1^HAC^ to directly compete with (PE-conjugated) durvalumab for binding to PDL1 on the surface of RKO cells. We discovered that sPD1^HAC^ effectively out-competed durvalumab (Figure 4C). These results demonstrate that sPD1^HAC^, regardless of the producer cell type, could significantly block the binding of the durvalumab antibody. In fact, MSC-produced sPD1**^HAC^** could reduce the binding of rhPD1 and the durvalumab signal to below 6%, demonstrating its potential superior therapeutic function (Figure 4B and 4C).

**Figure 4.**
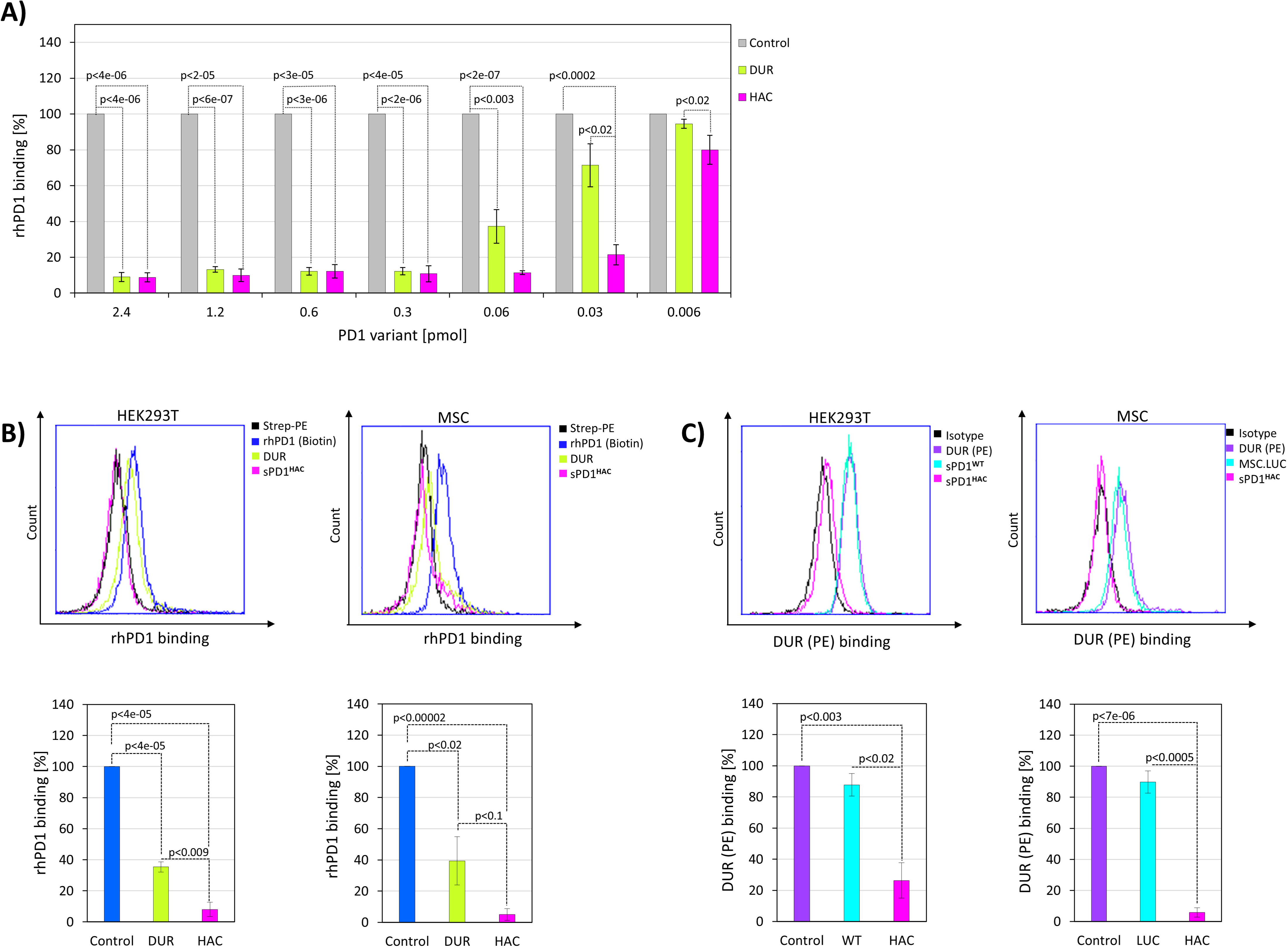
MSC-produced sPD1HAC (MSC.sPD1HAC) outcompetes durvalumab. **A** Wells of a 96-well plate were coated with rhPDL1, incubated with rhPD1Biotin, followed by Streptavidin-HRP and HRP substrate. To compare the ability of sPD1^HAC^ produced by HEK293T cells with equimolar amounts of durvalumab to compete for PD-L1 binding, various concentrations (0.006–2.4 pmol) of durvalumab (green) or sPD1^HAC^ (pink) were added. Controls using equivalent volumes of HEK293T supernatant are shown in grey, with their values set to 100. **B** Flow cytometric analysis of RKO cells stained with rhPD1Biotin and Streptavidin-PE (blue), Streptavidin-PE-alone as a negative control (black), or rhPD1Biotin and Streptavidin-PE in the presence of either sPD1^HAC^ (pink) or durvalumab-(green). Representative FACS plots are shown in the upper panel, with quantitative analyses in the lower panel. sPD1^HAC^ was produced in HEK293T cells (left) or MSCs (right). **C** Flow cytometric analysis of RKO cells stained with durvalumab-PE (DUR (PE); violet), PE-conjugated isotype control (black), or durvalumab-PE in the presence of either sPD1^HAC^ (pink), an equivalent volume of sPD1^WT^ supernatant (left; turquoise), or an equivalent volume of MSC.Luc supernatant (right; turquoise). Representative FACS plots are shown in the upper panels, and quantitative analyses are shown in the lower panels. sPD1^HAC^ was produced in HEK293T cells (left) or MSCs (right). Data represent mean ± SE

### MSC-produced sPD1^HAC^ outcompetes mPD1 binding to mPDL1

Having analysed sPD1**^HAC^**with human PDL1 protein and human cells, we wondered whether it would also work with murine PDL1 (mPDL1), which is the prerequisite for MSC.sPD1^HAC^ efficacy testing in an *in vivo* mouse model. First, we transfected HEK293T cells with an mPDL1 expression construct and stained the resulting cells with three different anti-mouse PDL1 antibodies (MIH6, MIH7 and 10F.9G2). Despite efforts to titrate and optimise the concentrations of all three antibodies for our flow cytometric experiments, the staining yielded two mPDL1 signal peaks (Figure 5A). These results indicate that overexpression leads to (at least) two different versions of PDL1 displayed on the cellular surface, possibly distinguished by varying degrees of post-translational modifications. Regarding the inhibitory role of sPD1HAC, we obtained different results depending on the antibody used. While sPD1^HAC^ was able to outcompete the binding of MIH7 and 10F.9G2 antibodies, no such effect was found with the MIH6 antibody (Figure 5B). These findings are very likely caused by the different epitopes recognised by the three monoclonal antibodies. In this context, MIH6 does not share its binding surface with sPD1^HAC^. Notwithstanding, the results demonstrate that sPD1^HAC^ can block the interaction between a specific antibody and mPDL1.

**Figure 5.**
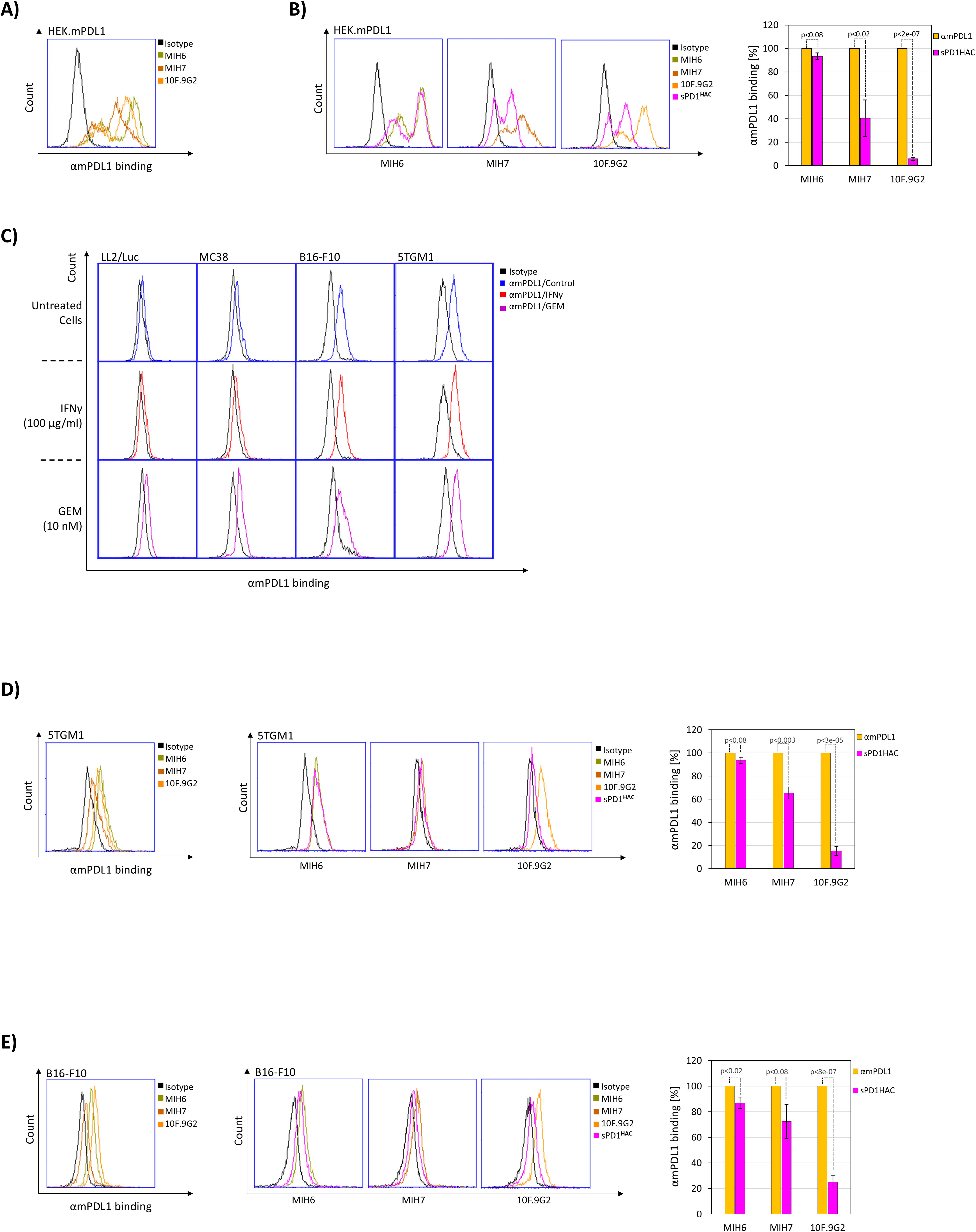
MSC-produced sPD1^HAC^ (MSC.sPD1^HAC^) outcompetes mouse anti-PDL1 antibodies for binding to murine PDL1 (mPDL1) **A** HEK293T cells were transfected with mouse PDL1 (mPDL1) and stained with different PE-conjugated anti-mPDL1 antibodies; 10F.9G2 (orange), MIH7 (brown), MIH6 (green), or the corresponding isotype (black). **B** mPDL1-overexpressing HEK293T cells were stained with PE-conjugated anti-mPDL1 clones MIH6 (green), MIH7 (brown), 10F.9G2 (orange), or the corresponding isotype control (black). In addition, each anti-mPDL1 antibody was tested in combination with sPD1^HAC^ (pink). Representative histograms are shown on the left, one for each mPDL1 antibody. Quantitative analyses of antibody binding (%) with or without sPD1^HAC^ are shown on the right (background-corrected-positive controls set to 100%). **C** Flow cytometric diagrams of four murine cancer cells (LL2/Luc, MC38, B16-F10 and 5TGM1) stained with an anti-mPDL1 antibody (10F.9G2). The cells were either left untreated or treated with IFN-γ (100 µg/ml) or Gemcitabine (GEM; 10 nM). **D** Flow cytometric analysis of 5TGM1 cells stained with PE-conjugated anti-mPDL1 antibodies MIH6 (green), MIH7 (brown), 10F.9G2 (orange), or the corresponding isotype control (black, left). Each anti-mPDL1 antibody was additionally tested in combination with sPD1^HAC^ (pink). Representative histograms of the competition assays are shown in the middle (one for each antibody), with quantitative analyses of antibody binding (%) with or without sPD1^HAC^ displayed on the right (background corrected-positive controls set to 100%). **E** Flow cytometric analysis of B16-F10 cells stained with PE-conjugated anti-mPDL1 antibodies MIH6 (green), MIH7 (brown), 10F.9G2 (orange), or the corresponding isotype control (black, left). The ability of sPD1^HAC^ to outcompete the respective anti-mPDL1 antibody was also evaluated (pink). Representative histograms of the competition assays are shown in the middle (one per antibody), while quantitative binding analyses (%) with or without sPD1^HAC^ are displayed on the right (background-corrected positive-controls set to 100%).

Next, we tested several murine cancer cell lines for their endogenous mPDL1 expression using the 10F.9G2 antibody and found the melanoma cell line B16-F10 and the multiple myeloma cell line 5TGM1 to be positive (Figure 5C). In contrast, the Lewis Lung carcinoma cell line LL2/Luc and the colorectal cancer cell line MC38 exhibited very low expression, which moderately increased after gemcitabine treatment but did not increase substantially after IFN-γ (Figure 5C). To test the inhibitory potential of MSC-produced sPD1**^HAC^**, we used 5TGM1 (Figure 5D) and B16-F10 (Figure 5E) cells with endogenous levels of PDL1. Similar to the findings with mPDL1-overexpressing HEK293T cells, we saw the inhibition was dependent on the anti-PDL1 antibody used. The binding of 10F.9G2 could be blocked almost completely by sPD1^HAC^, whereas the binding of MIH7 could be only slightly decreased, and MIH6 binding was hardly affected (Figure 5D and 5E). Overall, the results indicate that MSC.sPD1^HAC^ warrants further testing in a murine cancer model.

### MSC.sPD1^HAC^ exhibits anti-tumour activity in a melanoma model

After having established the ability of sPD1^HAC^ to inhibit the binding of PD1 and PDL1, and even the docking of specific antibodies to PDL1, in both human and murine systems, we went on to test the ICI and anti-cancer functions in a mouse model. For this, we used B16-F10 melanoma cells that form tumour lesions in the lungs of mice within a short period of time and have frequently been used in the study of ICIs (47–49). The B16-F10 cells were injected intravenously into C57BL/6 mice, and seven days later, we administered MSC.sPD1^HAC^ and MSC.LUC control cells. After another seven days, the animals were euthanised and their lungs histologically examined for lesions (Figure 6A). We found that treatment with MSC.sPD1^HAC^ resulted in a marked reduction in the macroscopic appearance (Figure 6B) and in the number and size of tumour nodules in the lungs (Figure 6C-E) of these mice. On average, MSC.sPD1^HAC^ treatment reduced the number of lung lesions by 5-fold. These results provide strong evidence for the therapeutic activity of MSC.sPD1^HAC^, as it even functions in the suboptimal context of a murine model for which the HAC variant was not designed.

**Figure 6.**
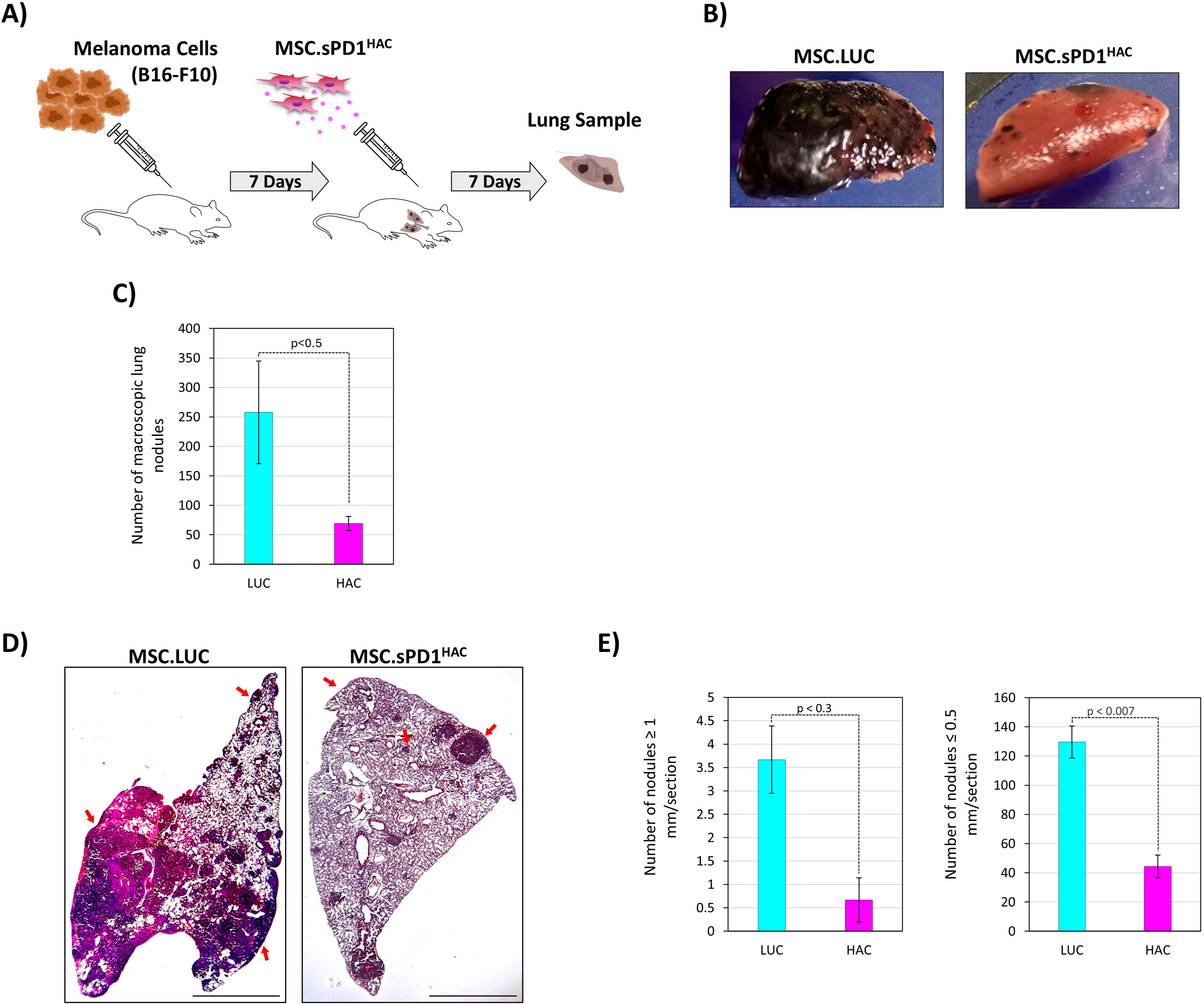
Therapeutic delivery of sPD1HAC by MSCs suppresses lung lesions in a B16-F10 melanoma mouse model. **A** Schematic overview of the experimental design. C57BL/6 mice were intravenously injected with 1×10^6^ B16-F10 cells, followed 7 days later by 1×10^6^ MSC.PD1^HAC^ (or MSC.LUC as controls). Another 7 days later, the lungs were harvested for subsequent analysis of tumour lesions. **B** Representative images from formalin-fixed lungs from tumour-bearing animals treated with MSC.LUC or MSC.sPD1^HAC^. **C** Quantification of macroscopic lung lesions from tumour-bearing mice treated with MSC.LUC (LUC) or MSC.sPD1^HAC^ (HAC). (n=7 animals/group) **D** H&E-stained lung sections from mice treated with MSC.LUC or MSC.sPD1^HAC^. Nodules are indicated by red arrows. Scale bar = 2800 µm. **E** Nodules from tumour-bearing mice treated with MSC.LUC (LUC) or MSC.sPD1^HAC^ (HAC) were quantified according to their size. For this, 54 fields/lung were analysed.

## Discussion

This study demonstrates the successful engineering and use of MSCs to secrete a high-affinity PD1 variant fused to IgG1-Fc, termed sPD1^HAC^, offering a novel strategy to target the PD1/PDL1 axis in cancer immunotherapy. Previous studies showed that this high-affinity HAC variant, developed by yeast surface display, achieved an up to 40,000-fold enhancement over wild-type PD1 (36). Furthermore, resolving the crystal structure of the HAC PD-1 protein bound to PDL1 revealed that the variant engages PDL1 through an expanded network of polar interactions. These are mediated by mutations such as M70E, Y68H, and K78T, which stabilise the interface via hydrogen bonds and salt bridges. An additional aspect of the HAC PD1 design is its pH-sensitive binding behaviour. The protonation-dependent interaction between H68 and D122 in PDL1 confers enhanced affinity under acidic conditions, mimicking the tumour microenvironment (TME). This property may enable selective targeting of tumour tissue while sparing normal cells (50).

While these characteristics of the HAC variant offer a good degree of tumour specificity, systemic ICI administration still faces several issues, including relatively poor tumour penetration, remaining off-target toxicity, and resistance mechanisms within the TME (33, 51). Therefore, we set out to combine the advantages of the HAC variant with the properties of MSC-mediated gene delivery (31, 33, 39). MSCs possess an innate ability to home to TMEs (52). This property is largely attributed to chemokine gradients, adhesion molecules, and inflammatory signals secreted by tumour tissues, which mimic wound healing cues and attract MSCs via receptors such as CXCR4 (53–56) This tumour tropism of MSCs enables spatially restricted ICI delivery, which in the future could be boosted by engineering MSC.sPD1^HAC^ to co-express the chemokine receptor CXCR4, Epidermal Growth Factor Receptor (EGFR) or artificial tumour-binding receptors (53, 57–60). Moreover, it was shown that hypoxic preconditioning or exposure to inflammatory cytokines prior to administration can improve migration of MSCs to hypoxic TMEs and enhance their responsiveness to tumour-emitted factors (61, 62). Additionally, biophysical techniques such as using MSCs labelled with magnetic nanoparticles allow external magnetic fields to enrich them at tumour sites (63). Thus, there are several approaches by which the tumour homing activities of MSC.sPD1^HAC^ can be further strengthened to potentially increase therapeutic functionality. Furthermore, MSCs are able to reduce side effects like irAEs and, for example, have been shown to prevent ICI-induced type 1 diabetes in murine models by suppressing CXCL9+ macrophage accumulation in pancreatic islets (64, 65). Therefore, MSC.sPD1^HAC^ therapy may exert a synergistic effect by directly targeting the tumour with sPD1^HAC^ while at the same time preventing some of the undesired irAEs via the immune-dampening activities of MSCs. However, this is a balancing act as on the other side ICIs reverse the MSC-mediated immune-suppression (66). Fortunately, the immune-modulatory characteristics of MSCs can be regulated by various priming protocols using Toll-like receptor signalling. There, MSCs are polarised into pro-inflammatory MSC1 by TLR4 stimulation or into an immunosuppressive MSC2 phenotype by TLR3 ligands (67, 68). Hence, the overall immune-regulatory status of MSC.sPD1^HAC^ can be fine-tuned using TLR priming to achieve the best balance for any given application and/or cancer.

In the present study, we tested MSCs expressing sPD1^HAC^ in the B16-F10 melanoma model. We were able to significantly reduce the tumour burden with a single intravenous injection of engineered MSCs, in contrast to the HAC recombinant protein, which had to be injected daily over two weeks (36). Melanoma was the first cancer in which ICIs were clinically approved and used with great success (69–71). Having shown proof-of-principle for this tumour type has set the stage to expand and test the utility of MSC.sPD1 ^HAC^ in other solid cancer models. MSC.sPD1^HAC^ could also be further engineered to turn non-inflamed (cold) tumours into T cell-inflamed (hot) phenotypes by co-expression of TNFSF14/LIGHT (72). LIGHT has been shown to be a potent inducer of T cell infiltration via lymphotoxin β-receptor signalling leading to increased CD4+ and CD8+ T cell infiltration of the tumour tissue. These findings suggest that MSCs co-expressing LIGHT and sPD1^HAC^ may synergistically overcome tumour immune resistance.

We assessed three types of MSCs (adipose tissue, bone marrow, and umbilical cord) for their utility in our ICI approach by analysing their PDL1 expression levels. AT-MSCs exhibited the lowest levels of PDL1 on their surface, making them the most suitable vehicle for delivering sPD1^HAC^, as they would not scavenge their own therapeutic payload. In the context of using tissue-derived MSCs in a cell therapeutic application, it is important to note that they are relatively easy to culture, but their limited lifespan and scalability pose challenges for clinical use. iPSC-derived MSCs (iMSCs) offer a potentially unlimited, autologous source, generated under defined conditions involving TGF-β inhibition and extracellular matrix support (73). However, their PDL1 surface expression needs to be tested, and their tumour tropism evaluated and compared before their use.

Together, these studies exemplify how protein engineering can overcome the limitations of antibody therapeutics by enhancing tissue accessibility, minimising off-target immune effects and achieving continuous expression and availability of the therapeutic agent. Future work should explore MSC.sPD1 in other cancer models and investigate combinatorial strategies with other immune modulators.

## Acknowledgments

This work was funded by a grant from Pancreatic Cancer UK (RMZ). PEB is funded by the East Suffolk and North Essex NHS Foundation Trust. SGD is funded by the Turkish Ministry of National Education.

## Author contribution

AM, SR, GNB and RMZ supervised the study; SG, NK, and PB performed the *in vitro* experiments and analysed data; AM and TC performed and analysed the *in vivo* experiments. SG, PB and TC created the figures. AM and RMZ wrote the paper; SG, PB, and GNB edited the manuscript; all authors read and reviewed the manuscript.

## References

1. Wei SC, Duffy CR, Allison JP. Fundamental Mechanisms of Immune Checkpoint Blockade Therapy. Cancer Discov. 2018;8(9):1069–86.

2. Mc Neil V, Lee SW. Advancing Cancer Treatment: A Review of Immune Checkpoint Inhibitors and Combination Strategies. Cancers (Basel). 2025;17(9).

3. Linsley PS, Brady W, Urnes M, Grosmaire LS, Damle NK, Ledbetter JA. CTLA-4 is a second receptor for the B cell activation antigen B7. J Exp Med. 1991;174(3):561–9.

4. Ishida Y, Agata Y, Shibahara K, Honjo T. Induced expression of PD-1, a novel member of the immunoglobulin gene superfamily, upon programmed cell death. Embo j. 1992;11(11):3887–95.

5. Dong H, Zhu G, Tamada K, Chen L. B7-H1, a third member of the B7 family, co-stimulates T-cell proliferation and interleukin-10 secretion. Nat Med. 1999;5(12):1365–9.

6. Freeman GJ, Long AJ, Iwai Y, Bourque K, Chernova T, Nishimura H, et al. Engagement of the PD-1 immunoinhibitory receptor by a novel B7 family member leads to negative regulation of lymphocyte activation. J Exp Med. 2000;192(7):1027–34.

7. Keir ME, Butte MJ, Freeman GJ, Sharpe AH. PD-1 and its ligands in tolerance and immunity. Annu Rev Immunol. 2008;26:677–704.

8. Rowshanravan B, Halliday N, Sansom DM. CTLA-4: a moving target in immunotherapy. Blood. 2018;131(1):58–67.

9. Sharma P, Goswami S, Raychaudhuri D, Siddiqui BA, Singh P, Nagarajan A, et al. Immune checkpoint therapy-current perspectives and future directions. Cell. 2023;186(8):1652–69.

10. Zamani MR, Šácha P. Immune checkpoint inhibitors in cancer therapy: what lies beyond monoclonal antibodies? Med Oncol. 2025;42(7):273.

11. Hodi FS, O’Day SJ, McDermott DF, Weber RW, Sosman JA, Haanen JB, et al. Improved survival with ipilimumab in patients with metastatic melanoma. N Engl J Med. 2010;363(8):711–23.

12. Topalian SL, Hodi FS, Brahmer JR, Gettinger SN, Smith DC, McDermott DF, et al. Safety, activity, and immune correlates of anti-PD-1 antibody in cancer. N Engl J Med. 2012;366(26):2443–54.

13. Yokosuka T, Takamatsu M, Kobayashi-Imanishi W, Hashimoto-Tane A, Azuma M, Saito T. Programmed cell death 1 forms negative costimulatory microclusters that directly inhibit T cell receptor signaling by recruiting phosphatase SHP2. J Exp Med. 2012;209(6):1201–17.

14. Sheppard KA, Fitz LJ, Lee JM, Benander C, George JA, Wooters J, et al. PD-1 inhibits T-cell receptor induced phosphorylation of the ZAP70/CD3zeta signalosome and downstream signaling to PKCtheta. FEBS Lett. 2004;574(1-3):37–41.

15. Nishimura H, Honjo T. PD-1: an inhibitory immunoreceptor involved in peripheral tolerance. Trends Immunol. 2001;22(5):265–8.

16. Atsaves V, Leventaki V, Rassidakis GZ, Claret FX. AP-1 Transcription Factors as Regulators of Immune Responses in Cancer. Cancers (Basel). 2019;11(7).

17. Arranz-Nicolás J, Martin-Salgado M, Adán-Barrientos I, Liébana R, Del Carmen Moreno-Ortíz M, Leitner J, et al. Diacylglycerol kinase α inhibition cooperates with PD-1-targeted therapies to restore the T cell activation program. Cancer Immunol Immunother. 2021;70(11):3277–89.

18. Jenkins E, Whitehead T, Fellermeyer M, Davis SJ, Sharma S. The current state and future of T-cell exhaustion research. Oxf Open Immunol. 2023;4(1):iqad006.

19. Ghiotto M, Gauthier L, Serriari N, Pastor S, Truneh A, Nunès JA, et al. PD-L1 and PD-L2 differ in their molecular mechanisms of interaction with PD-1. Int Immunol. 2010;22(8):651–60.

20. Wang JC, Sun L. PD-1/PD-L1, MDSC Pathways, and Checkpoint Inhibitor Therapy in Ph(-) Myeloproliferative Neoplasm: A Review. Int J Mol Sci. 2022;23(10).

21. Beenen AC, Sauerer T, Schaft N, Dorrie J. Beyond Cancer: Regulation and Function of PD-L1 in Health and Immune-Related Diseases. Int J Mol Sci. 2022;23(15).

22. Mazanet MM, Hughes CC. B7-H1 is expressed by human endothelial cells and suppresses T cell cytokine synthesis. J Immunol. 2002;169(7):3581–8.

23. Swallow MM, Wallin JJ, Sha WC. B7h, a novel costimulatory homolog of B7.1 and B7.2, is induced by TNFalpha. Immunity. 1999;11(4):423–32.

24. Lomphithak T, Duangthim N, Sonkaew S, Jitkaew S. Necroptosis-driven T cell activation promotes IL-6-mediated PD-L1 upregulation in cholangiocarcinoma cells: IL-6 gene signature as a biomarker for chemo-immunotherapy response. Biol Direct. 2025;20(1):98.

25. Gorchs L, Fernandez Moro C, Bankhead P, Kern KP, Sadeak I, Meng Q, et al. Human Pancreatic Carcinoma-Associated Fibroblasts Promote Expression of Co-inhibitory Markers on CD4(+) and CD8(+) T-Cells. Front Immunol. 2019;10:847.

26. Iwai Y, Ishida M, Tanaka Y, Okazaki T, Honjo T, Minato N. Involvement of PD-L1 on tumor cells in the escape from host immune system and tumor immunotherapy by PD-L1 blockade. Proc Natl Acad Sci U S A. 2002;99(19):12293–7.

27. Sharpe AH, Pauken KE. The diverse functions of the PD1 inhibitory pathway. Nat Rev Immunol. 2018;18(3):153–67.

28. Bayless NL, Bluestone JA, Bucktrout S, Butterfield LH, Jaffee EM, Koch CA, et al. Development of preclinical and clinical models for immune-related adverse events following checkpoint immunotherapy: a perspective from SITC and AACR. J Immunother Cancer. 2021;9(9).

29. Abraham PE, Johnson DB. Long-Term Toxicities of Immune Checkpoint Inhibitors. Drugs. 2025;85(12):1535–49.

30. Wang Y, Abu-Sbeih H, Mao E, Ali N, Ali FS, Qiao W, et al. Immune-checkpoint inhibitor-induced diarrhea and colitis in patients with advanced malignancies: retrospective review at MD Anderson. J Immunother Cancer. 2018;6(1):37.

31. Mohr A, Albarenque SM, Deedigan L, Yu R, Reidy M, Fulda S, et al. Targeting of XIAP Combined with Systemic Mesenchymal Stem Cell-Mediated Delivery of sTRAIL Ligand Inhibits Metastatic Growth of Pancreatic Carcinoma Cells. Stem Cells. 2010;28(11):2109–20.

32. Mohr A, Chu T, Brooke GN, Zwacka RM. MSC.sTRAIL Has Better Efficacy than MSC.FL-TRAIL and in Combination with AKTi Blocks Pro-Metastatic Cytokine Production in Prostate Cancer Cells. Cancers (Basel). 2019;11(4):568.

33. Albarenque SM, Zwacka RM, Mohr A. Both human and mouse mesenchymal stem cells promote breast cancer metastasis. Stem Cell Res. 2011;7(2):163–71.

34. Mohr A, Chu T, Clarkson CT, Brooke GN, Teif VB, Zwacka RM. Fas-threshold signalling in MSCs promotes pancreatic cancer progression and metastasis. Cancer Lett. 2021;519:63–77.

35. Studeny M, Marini FC, Champlin RE, Zompetta C, Fidler IJ, Andreeff M. Bone marrow-derived mesenchymal stem cells as vehicles for interferon-beta delivery into tumors. Cancer Res. 2002;62(13):3603–8.

36. Maute RL, Gordon SR, Mayer AT, McCracken MN, Natarajan A, Ring NG, et al. Engineering high-affinity PD-1 variants for optimized immunotherapy and immuno-PET imaging. Proc Natl Acad Sci U S A. 2015;112(47):E6506–14.

37. Yu R, Albarenque SM, Cool RH, Quax WJ, Mohr A, Zwacka RM. DR4 specific TRAIL variants are more efficacious than wild-type TRAIL in pancreatic cancer. Cancer Biol Ther. 2014;15(12):1658–66.

38. Yu R, Deedigan L, Albarenque SM, Mohr A, Zwacka RM. Delivery of sTRAIL variants by MSCs in combination with cytotoxic drug treatment leads to p53-independent enhanced antitumor effects. Cell Death and Disease. 2013;4:e503.

39. Mohr A, Lyons M, Deedigan L, Harte T, Shaw G, Howard L, et al. Mesenchymal stem cells expressing TRAIL lead to tumour growth inhibition in an experimental lung cancer model. Journal of Cellular and Molecular Medicine. 2008;12(6B):2628–43.

40. Baldini C, Champiat S, Vuagnat P, Massard C. durvalumab for the management of urothelial carcinoma: a short review on the emerging data and therapeutic potential. Onco Targets Ther. 2019;12:2505–12.

41. Cheng Y, Spigel DR, Cho BC, Laktionov KK, Fang J, Chen Y, et al. durvalumab after Chemoradiotherapy in Limited-Stage Small-Cell Lung Cancer. N Engl J Med. 2024;391(14):1313–27.

42. Antonia SJ, Villegas A, Daniel D, Vicente D, Murakami S, Hui R, et al. durvalumab after Chemoradiotherapy in Stage III Non-Small-Cell Lung Cancer. N Engl J Med. 2017;377(20):1919–29.

43. Powles T, Catto JWF, Galsky MD, Al-Ahmadie H, Meeks JJ, Nishiyama H, et al. Perioperative durvalumab with Neoadjuvant Chemotherapy in Operable Bladder Cancer. N Engl J Med. 2024;391(19):1773–86.

44. Oh DY, He AR, Qin S, Chen LT, Okusaka T, Kim JW, et al. durvalumab plus chemotherapy in advanced biliary tract cancer: 3-year overall survival update from the phase III TOPAZ-1 study. J Hepatol. 2025;83(5):1092–101.

45. Aggarwal C, Martinez-Marti A, Majem M, Barlesi F, Carcereny E, Chu Q, et al. durvalumab Alone or Combined With Novel Agents for Unresectable Stage III Non-Small Cell Lung Cancer: Update From the COAST Randomized Clinical Trial. JAMA Netw Open. 2025;8(7):e2518440.

46. Park J, Skålhegg BS. Combination of PD-1/PD-L1 and CTLA-4 inhibitors in the treatment of cancer - a brief update. Front Immunol. 2025;16:1680838.

47. Klement JD, Redd PS, Lu C, Merting AD, Poschel DB, Yang D, et al. Tumor PD-L1 engages myeloid PD-1 to suppress type I interferon to impair cytotoxic T lymphocyte recruitment. Cancer Cell. 2023;41(3):620–36.e9.

48. Lin H, Wei S, Hurt EM, Green MD, Zhao L, Vatan L, et al. Host expression of PD-L1 determines efficacy of PD-L1 pathway blockade-mediated tumor regression. J Clin Invest. 2018;128(2):805–15.

49. Vijayakumar G, McCroskery S, Palese P. Engineering Newcastle Disease Virus as an Oncolytic Vector for Intratumoral Delivery of Immune Checkpoint Inhibitors and Immunocytokines. J Virol. 2020;94(3).

50. Pascolutti R, Sun X, Kao J, Maute RL, Ring AM, Bowman GR, et al. Structure and Dynamics of PD-L1 and an Ultra-High-Affinity PD-1 Receptor Mutant. Structure. 2016;24(10):1719–28.

51. Alsaafeen BH, Ali BR, Elkord E. Resistance mechanisms to immune checkpoint inhibitors: updated insights. Mol Cancer. 2025;24(1):20.

52. Mohr A, Zwacka R. The future of mesenchymal stem cell-based therapeutic approaches for cancer - From cells to ghosts. Cancer Lett. 2018;414:239–49.

53. Yang JX, Zhang N, Wang HW, Gao P, Yang QP, Wen QP. CXCR4 receptor overexpression in mesenchymal stem cells facilitates treatment of acute lung injury in rats. J Biol Chem. 2015;290(4):1994–2006.

54. Dvorak HF. Tumors: wounds that do not heal-redux. Cancer Immunol Res. 2015;3(1):1–11.

55. Li P, Gong Z, Shultz LD, Ren G. Mesenchymal stem cells: From regeneration to cancer. Pharmacol Ther. 2019;200:42–54.

56. Jiang D, Scharffetter-Kochanek K. Mesenchymal Stem Cells Adaptively Respond to Environmental Cues Thereby Improving Granulation Tissue Formation and Wound Healing. Front Cell Dev Biol. 2020;8:697.

57. Kalimuthu S, Oh JM, Gangadaran P, Zhu L, Lee HW, Rajendran RL, et al. In Vivo Tracking of Chemokine Receptor CXCR4-Engineered Mesenchymal Stem Cell Migration by Optical Molecular Imaging. Stem Cells Int. 2017;2017:8085637.

58. Park SA, Ryu CH, Kim SM, Lim JY, Park SI, Jeong CH, et al. CXCR4-transfected human umbilical cord blood-derived mesenchymal stem cells exhibit enhanced migratory capacity toward gliomas. Int J Oncol. 2011;38(1):97–103.

59. Sato H, Kuwashima N, Sakaida T, Hatano M, Dusak JE, Fellows-Mayle WK, et al. Epidermal growth factor receptor-transfected bone marrow stromal cells exhibit enhanced migratory response and therapeutic potential against murine brain tumors. Cancer Gene Ther. 2005;12(9):757–68.

60. Komarova S, Roth J, Alvarez R, Curiel DT, Pereboeva L. Targeting of mesenchymal stem cells to ovarian tumors via an artificial receptor. J Ovarian Res. 2010;3:12.

61. Lee B-C, Kang K-S. Functional enhancement strategies for immunomodulation of mesenchymal stem cells and their therapeutic application. Stem Cell Research & Therapy. 2020;11(1):397.

62. Xinaris C, Morigi M, Benedetti V, Imberti B, Fabricio AS, Squarcina E, et al. A novel strategy to enhance mesenchymal stem cell migration capacity and promote tissue repair in an injury specific fashion. Cell Transplant. 2013;22(3):423–36.

63. Gundersen RA, Chu T, Abolfathi K, Dogan SG, Blair PE, Nago N, et al. Generation of magnetic biohybrid microrobots based on MSC.sTRAIL for targeted stem cell delivery and treatment of cancer. Cancer Nanotechnol. 2023;14:54.

64. Hu H, Zakharov PN, Peterson OJ, Unanue ER. Cytocidal macrophages in symbiosis with CD4 and CD8 T cells cause acute diabetes following checkpoint blockade of PD-1 in NOD mice. Proc Natl Acad Sci U S A. 2020;117(49):31319–30.

65. Kawada-Horitani E, Kita S, Okita T, Nakamura Y, Nishida H, Honma Y, et al. Human adipose-derived mesenchymal stem cells prevent type 1 diabetes induced by immune checkpoint blockade. Diabetologia. 2022;65(7):1185–97.

66. Kamrani S, Naseramini R, Khani P, Razavi ZS, Afkhami H, Atashzar MR, et al. Mesenchymal stromal cells in bone marrow niche of patients with multiple myeloma: a double-edged sword. Cancer Cell International. 2025;25(1):117.

67. Bernardo ME, Fibbe WE. Mesenchymal stromal cells: sensors and switchers of inflammation. Cell Stem Cell. 2013;13(4):392–402.

68. Waterman RS, Tomchuck SL, Henkle SL, Betancourt AM. A new mesenchymal stem cell (MSC) paradigm: polarization into a pro-inflammatory MSC1 or an Immunosuppressive MSC2 phenotype. PLoS One. 2010;5(4):e10088.

69. Larkin J, Chiarion-Sileni V, Gonzalez R, Grob JJ, Rutkowski P, Lao CD, et al. Five-Year Survival with Combined Nivolumab and Ipilimumab in Advanced Melanoma. N Engl J Med. 2019;381(16):1535–46.

70. Robert C, Schachter J, Long GV, Arance A, Grob JJ, Mortier L, et al. Pembrolizumab versus Ipilimumab in Advanced Melanoma. N Engl J Med. 2015;372(26):2521–32.

71. Robert C. A decade of immune-checkpoint inhibitors in cancer therapy. Nature Communications. 2020;11(1):3801.

72. Skeate JG, Otsmaa ME, Prins R, Fernandez DJ, Da Silva DM, Kast WM. TNFSF14: LIGHTing the Way for Effective Cancer Immunotherapy. Frontiers in Immunology. 2020;Volume 11 - 2020.

73. Sabapathy V, Kumar S. hiPSC-derived iMSCs: NextGen MSCs as an advanced therapeutically active cell resource for regenerative medicine. J Cell Mol Med. 2016;20(8):1571–88.

