## Supplementary figures and images for "MSC-delivered High-Affinity Variants of soluble PD1 lead to tumour regression"

### Gokcen_Supplemental Figure 1_bioRxiv_2025.tif

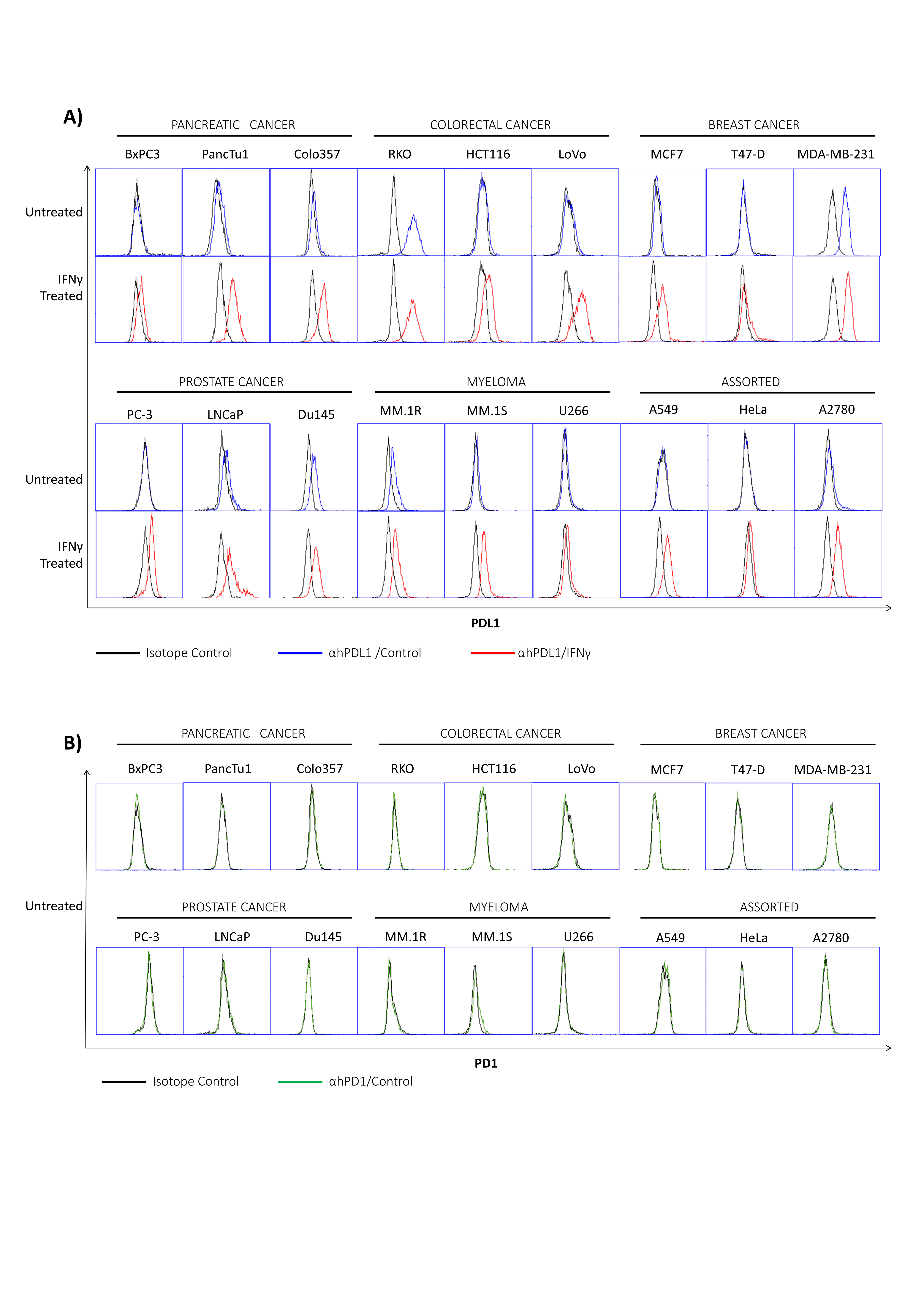
